# New observations on non-coding RNAs involved in the dual translation system in zebrafish development

**DOI:** 10.1101/869651

**Authors:** Timo M. Breit, Johanna F. B. Pagano, Pjotr L. van der Jagt, Ellis Mittring, Wim A. Ensink, Marina van Olst, Selina van Leeuwen, Wim de Leeuw, Ulrike Nehrdich, Herman P. Spaink, Han Rauwerda, Rob J. Dekker

## Abstract

Cellular translation relies heavily on the involvements of several types of non-coding RNAs. In previous studies we have identified a dual translation system in zebrafish development, involving maternal-type and somatic-type rRNAs, snoRNAs, and snRNAs. In this study we focused on several remaining non-coding RNAs involved in the translation system; tRNAs, RNase P, and SRP RNA. Even though our studies have been limited in extent, for all three types of non-coding RNA we were able to identify a maternal-specific type, with substantial sequence differences as compared to the somatic-type variant. Hence, these RNA types complement the previously discovered RNA types in the unique dual translation system in zebrafish development.

## Introduction

The oocyte is an intriguing and unique cell type for which the translation system functions quite differently as compared to somatic cells. Whereas in somatic cells translation generally quickly follows transcription, in oocytes mRNA is accumulated for later use in (early) embryogenesis. During this waiting period, these maternal mRNAs need to be silenced in order to maintain a normal cellular metabolism in oocytes (Tadros and Lipshitz 2005). Besides the production of mRNA from a specific set of maternal genes, mRNAs in mature oocytes are also atypical as they have no, or very short poly(A) tails (Graindorge et al. 2006; Weill et al. 2012; Subtelny et al. 2014), which likely play a role in the mRNA storage process.

Another distinction for oocyte RNA was discovered decades ago in Xenopus, where the expressed variant of 5S rRNA in oocytes differs from that in somatic cells. This maternal-type 5S RNA variant is gradually replaced throughout embryonic development by a somatic-type variant (Wegnez et al. 1972; Brown et al. 1977; Wormington and Brown 1983; Guinta et al. 1986; Komiya et al. 1986). Starting from this long-standing observation, our continuous studies reported in several manuscripts (Locati et al. 2017a, 2017b, 2018; Pagano et al. 2019b, 2019a) show the existence of a specialized maternal-type translation system in zebrafish far beyond just 5S rRNA. One by one, we have revealed that also the translation-involved non-coding RNAs 5.8S, 18S, 28S, snoRNA, and snRNA are implicated in this maternal-type translation system (Figure 1 and Table 1). The translation system however, has several other non-coding RNA components (Figure 1), i.e. miRNAs, tRNAs, RNase P, and SRP RNA. Here we report on several additional observations with respect to the latter non-coding RNAs, with the exception of miRNA, and their possible involvement in the maternal-specific translation system in zebrafish.

**Figure 1.**
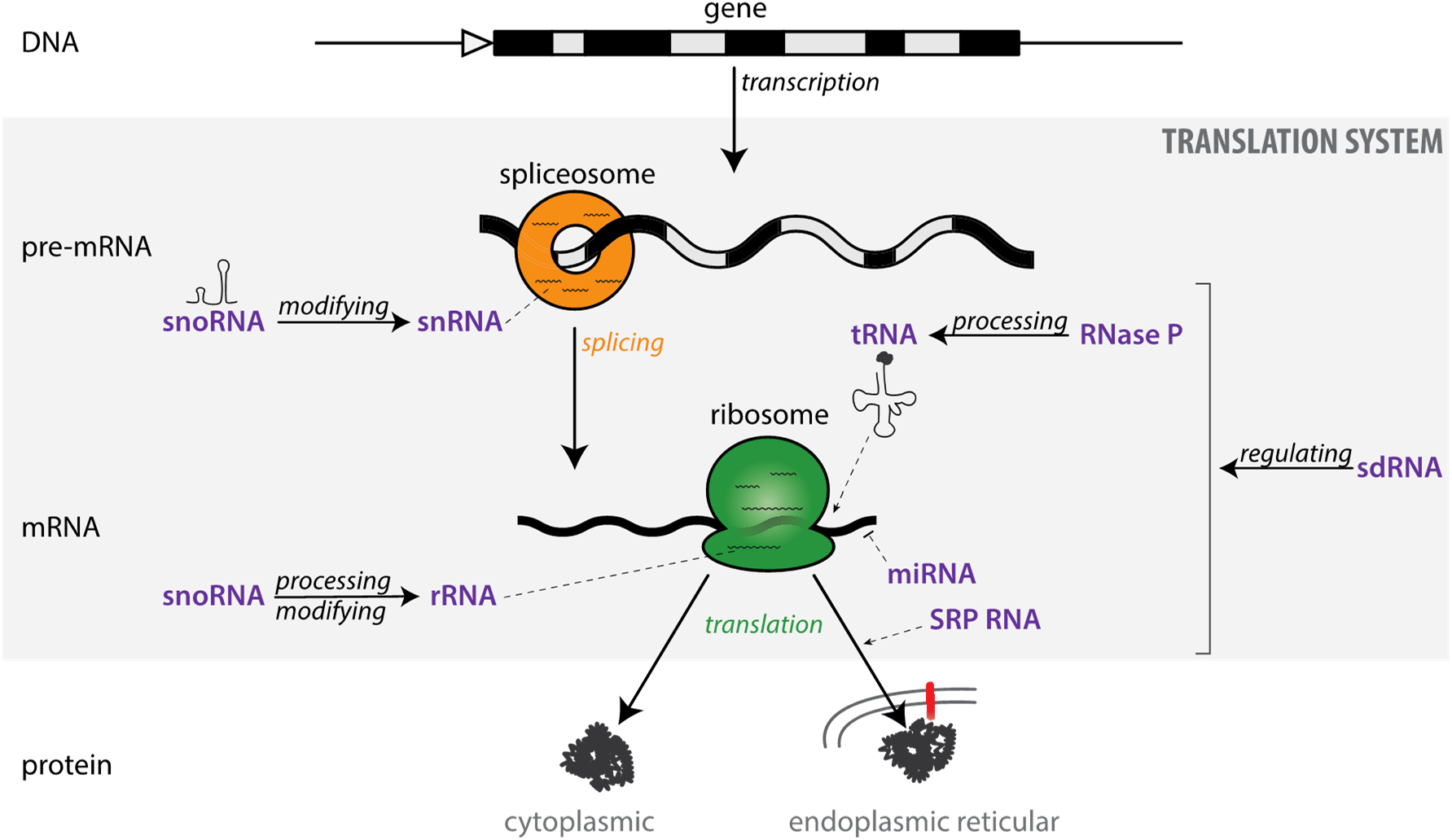
The translation system in the context of the central dogma. Schematic representation of the translation system in the central dogma of molecular biology, with a focus on the main involved non coding RNA types (purple)

**Table 1.** Main RNA types involved in the translation system.

| RNA processing & modifying | Sub-member | Size (~) | RNA polymerase | Primary target | Main process |
| --- | --- | --- | --- | --- | --- |
| snoRNA | H/ACA box | 91 nt | Pol II | RNA | RNA pseudouridylation |
|  | C/D box | 141 nt | Pol II | RNA | RNA methylation |
|  | U3 | 215 nt | Pol II or III | 45S | cleaving |
|  | U8 | 131 nt | Pol II or III | 45S | cleaving |
|  | snoZ30 | 104 nt | Pol II | U6 | RNA methylation |
| snRNA | U1 | 164 nt | Pol II | pre-mRNA | splicing |
|  | U2 | 191 nt | Pol II | pre-mRNA | splicing |
|  | U4 | 141 nt | Pol II | pre-mRNA | splicing |
|  | U5 | 118 nt | Pol II | pre-mRNA | splicing |
|  | U6 | 107 nt | Pol III | pre-mRNA | splicing |
| RNase P |  | 300 nt | Pol III | tRNA, lncRNA, rRNA, mRNA | cleaving |
| **Primary translation process** |  |  |  |  |  |
| rRNA | 5S | 120 nt | Pol III | mRNA | translation |
|  | 5.8S | 160 nt | Pol I | mRNA | translation |
|  | 18S | 1.900 nt | Pol I | mRNA | translation |
|  | 28S | 4.200 nt | Pol I | mRNA | translation |
| tRNA | 61 anticodons | 80 nt | Pol III | mRNA | translation |
| **Coding** |  |  |  |  |  |
| mRNA | many | variable | Pol II |  | translation |
| **Regulating RNA** |  |  |  |  |  |
| sdRNA | tRNA halves | 40 nt |  | retrotransposons, DNA | DNA processing |
| miRNA | many | 22 nt | Pol II | mRNA | degradation |
| **Auxiliary** |  |  |  |  |  |
| SRP RNA |  | 297 nt | Pol III | signal sequence of forming protein | translocating |

The here presented observations originate from initial analyses and preliminary results of available data and/or new experiments. Research funding constraints only allowed for limited investigation of these translation-involved non-coding RNA types. However, since these observations nicely complete the discovery of a dual translation system in zebrafish, we felt that this warranted sharing our preliminary findings on the three “missing” non-coding RNA types in this manuscript.

## tRNAs

### Background

Transfer RNAs (tRNAs) are central in the physical translation of the genetic code of mRNA into proteins in the context of ribosomes (Rich and RajBhandary 1976; Rodnina and Wintermeyer 2011). Yet, tRNAs are increasingly recognized as important players in a wide range of cellular processes (Schimmel 2017; Schorn and Martienssen 2018). tRNA biogenesis encompasses: translation of the precursor-tRNA, removal of the 5’ leader sequence, trimming of the 3’ trailer sequence, addition of CAA nucleotides, splicing-out of possible intronic sequences, and numerous nucleotide modifications (Phizicky and Hopper 2010; Kirchner and Ignatova 2015). Mature tRNAs are about 80 nucleotides long and have a characteristic cloverleaf secondary structure, as well as a L-shaped tertiary structure (Rich and RajBhandary 1976). In eukaryotes there are 61 possible anticodons and thus, in principle, 61 possible tRNAs (Alberts et al. 2015). However, due to the “Wobble principle”, which allows an amino acid to be coded by a set of anticodons (n = 2, 4, or 6), named isoacceptors (Goodenbour and Pan 2006) via effectively ‘ignoring’ the third base of the codon, a cell does not need all 61 tRNAs for proper translation. The minimum number of (anti)codons is 31, and although the exact number is different for each species, in general about 44 different anticodon tRNA genes are present in the genome of a species (Marck and Grosjean 2002). At the same time, each anticodon tRNA is usually present by multiple copies in the genome and if these copies of one tRNA anticodon have differences in their body sequence, they are called isodecoders (Goodenbour and Pan 2006).

One of the reasons for many isodecoders is the possibility to use different sets of isodecoders for each tRNA anticodon for different cell types, for instance in the case of pathologies (Rogers et al. 2010; Gingold et al. 2014; Kirchner and Ignatova 2015). Differential tRNA expression in combination with codon bias (Quax et al. 2015) and in relation to protein folding (Marín et al. 2017) is currently an emerging field of research.

### Zebrafish tRNA genomic organization

Given the existence of a maternal-specific ribosome in zebrafish development (Locati et al. 2017a, 2017b), the question was whether, similarly, a maternal-specific tRNA repertoire is used in early embryogenesis. To address this, an inventory was made (cf. M&M (Locati et al. 2017a)) of the tRNA genes starting from the genomic tRNA database GtRNAdb (release 2010) (Chan and Lowe 2009), which predicts a staggering 9,904 tRNA genes in the zebrafish genome (Zv9).

The first notable observation was the fact that the zebrafish genome contains all 61 types of anticodon tRNA (Supplemental Table ST1). Thus far, we have been unable to find any other species that has all the anticodon tRNAs present in its genome. For each anticodon, several tRNA genes are present in the zebrafish genome, ranging from minimal one (Asp-ATC) gene to maximal 825 (Asn-GTT) identical copies, as well as isodecoders. Furthermore, anticodon tRNAs appear to be organized in large clusters, small clusters, and/or solitary genes (Figure 2A, Supplemental Table ST1). The largest clusters per anticodon tRNA are primarily (~56%) found on the right arm of chromosome 4, but are also present on several other chromosomes (Figure 2A). Conversely, for each group of isoacceptors, one anticodon tRNA does not appear in a big gene cluster (n>16) (Supplemental Table ST1). These are the anticodon tRNA genes that are also often absent in genomes of other species (Marck and Grosjean 2002). The anticodons of these low-presence tRNA genes distribute evenly over the genetic code (Figure 2B) and involve a C or U base at their 3^rd^ position, which is in line with the possible presence of inosine in the tRNA anticodon (Agris et al. 2018).

**Figure 2.**
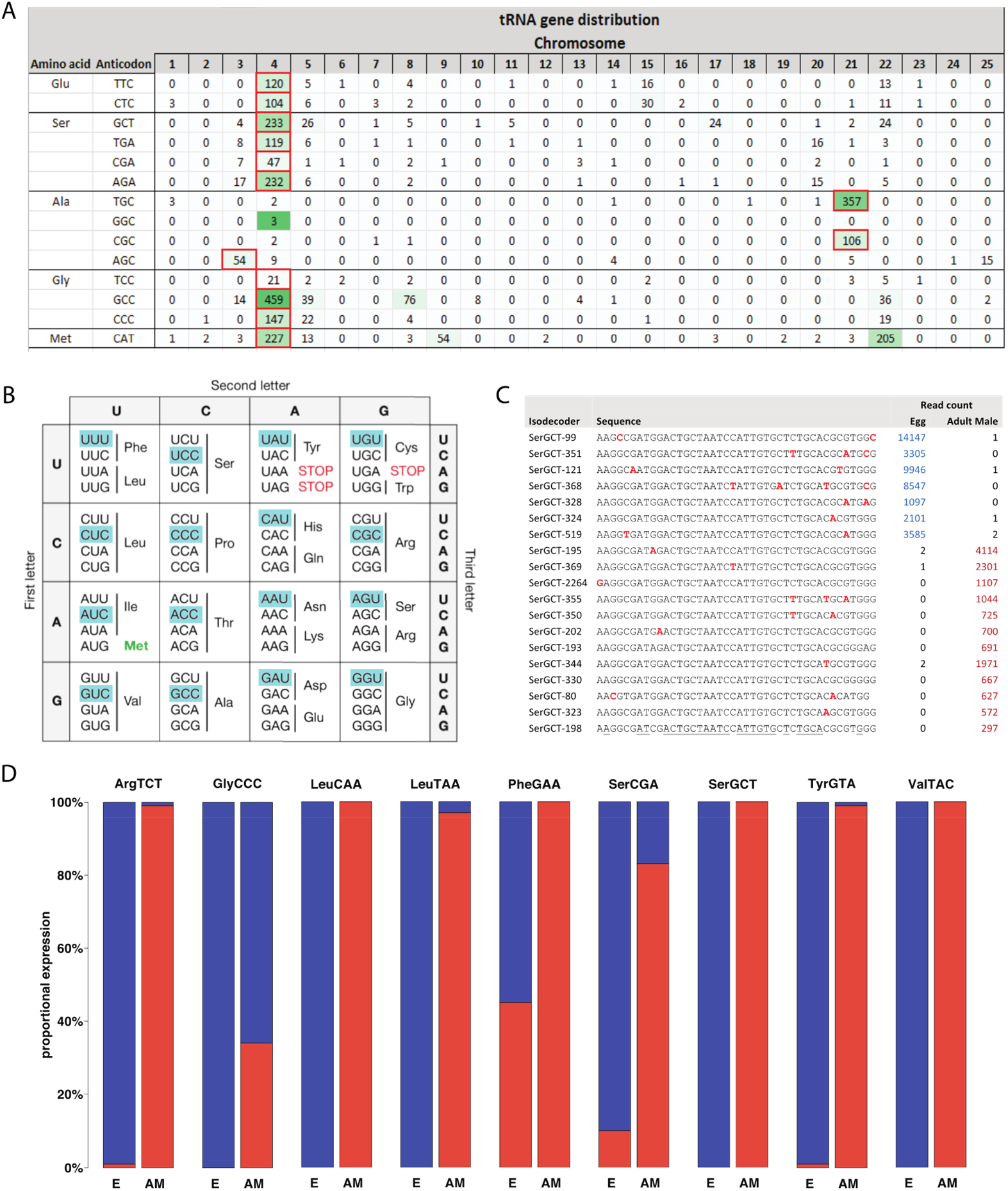
Maternal-type tRNA variant characteristics. A: The genomic distribution of some exemplary zebrafish tRNAs (complete table in Supplemental Table ST1). B: The genetic code table. For the highlighted codons, the associated anticodons tRNA genes are present in the genome with just a few copies. C: Alignment of exemplary tRNA SerGCT isodecoders and their differential expression in egg and adult male zebrafish. D: A barplot showing the proportional, average expression of maternal-type (blue) and somatic-type (red) variants for some exemplary anticodon tRNAs in egg (E) and adult male (AM) zebrafish (Supplemental Table ST1).

### Zebrafish tRNA maternal expression

As with 5S rRNA and snRNA, tRNAs are quite difficult to sequence by standard next-generation sequencing approaches, likely due to their strong secondary structure, 3’ amino acid, and many RNA modifications (Zheng et al. 2015). We therefore employed our RT-PCR-qSeq approach (cf. M&M (Pagano et al. 2019a)) with PCR primer pairs hybridizing to the 20 most 5’ terminal and 3’ terminal bases of nearly all anticodon tRNAs (Supplemental Table ST1). Regrettably, this approach leaves only about 40 nucleotides for isodecoder identification. For almost all investigated anticodon tRNAs, expression data was obtained by analyzing egg and adult male tail samples (cf. M&M (Pagano et al. 2019a)). An initial analysis clearly showed that there is differential expression between different isodecoders of each analyzed anticodon tRNA (Figure 2C and 2D). There are undoubtedly tRNA genes per anticodon which transcripts are either present in egg or adult tail. This demonstrates that also maternal-type tRNA genes exist. At the same time, we generally could not find obvious sequence signatures for maternal-type tRNAs. This however could be caused by the fact that we just examined the core of about 40 nucleotides for each tRNA gene, whereas there are often differences between isodecoders within the remaining parts of their tRNA body sequence, which are used in our method for PCR-primer binding, the 5’ and 3’ precursor tRNA parts, and/or intronic sequences of precursor tRNA.

What did stand out was that it seems as if the maternal-type tRNA genes are organized in large clusters, mostly on chromosome 4, whereas somatic-type genes are mostly found as single gene or in a small cluster. Yet, although the low-presence tRNA isoacceptor genes (Figure 2B) are organized like somatic-type genes, their transcripts actually are primarily present in egg (Supplemental Table ST1).

### Conclusion

Within the set of genes belonging to an anticodon tRNA, there are maternal-type and somatic-type isodecoders. The maternal-type isodecoders are mainly organized in large clusters on chromosome 4, which also harbor the maternal-type rRNA genes, yet without obvious sequence signatures. It would be interesting to determine whether the maternal-type and somatic-type tRNA genes line up with the reported “core” and “peripheral” tRNA genes proposed in *Drosophila* (Rogers et al. 2010). Also, maternal-type tRNAs might uniquely encompass all possible anticodons (n=61), which raises the question whether the zebrafish maternal-type ribosome might not allow for Wobble translation. An intriguing link might also be found in the codon identity that is involved in mRNA stability regulation and translation efficiency during the maternal-to-zygotic transition (Bazzini et al. 2016). Our tRNA observations must of course be confirmed by comprehensive experiments and analyses, but given these compelling preliminary observations, we postulate that tRNAs are a contributing part of the maternal-specific translation system.

## RNase P

### Background

Another type of ribonucleoprotein (RNP) involved in the cellular translation system is the RNase P enzyme. In vertebrates the RNase P enzyme consists of a non-coding RNA subunit (rpph1) and several proteins (reviewed in Klemm et al. 2016). Initially discovered for its role as a ribozyme that removes the 5’ leader of a precursor tRNA in the maturation process, it now is recognized to be also involved in rRNA and mRNA cleavage, as well as chromatin remodeling (reviewed in Jarrous 2017; Esakova and Krasilnikov 2010).

### Zebrafish RNase P genomic organization

Given the role of the RNase P enzyme in tRNA maturation and rRNA processing, the question was whether a maternal-specific RNase P enzyme exists in zebrafish. For this we focused on the non-coding RNA subunit of the RNase P enzyme. A rpph1 gene (Ribonuclease P RNA component H1) of 307 nucleotides is annotated in the zebrafish genome on chromosome 2 (position 37,826,734). A standard BLAST analysis (Madden 2002) revealed a similar (96%) gene copy (rpph1-sim), as well as a dissimilar (80%) gene copy (rpph1-dis), ~10 kb and ~5 Mb downstream respectively (Figure 3 A, Supplemental Table ST2).

**Figure 3:**
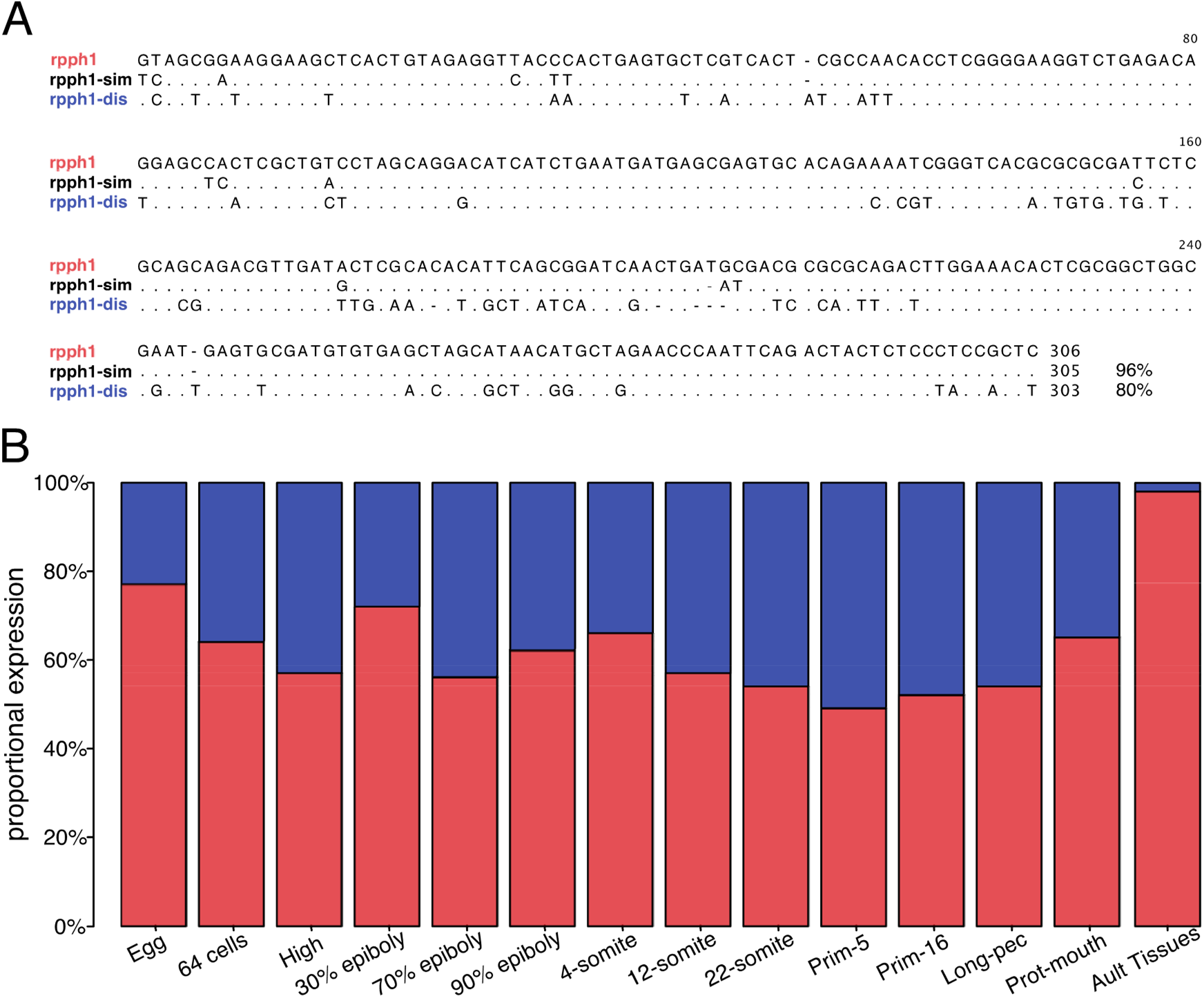
Maternal-type and somatic-type RNase P RNP variants. **A:** Sequence alignment of the three zebrafish RNase P RNP (rpph1) gene variants. Identical nucleotides or gaps are indicated as dots and dashes, respectively. **B:** Bar plot showing the maternal-type (blue) and somatic-type (red) rpph1 proportional expression during zebrafish developmental stages (Supplemental Table ST2).

**Figure 4.**
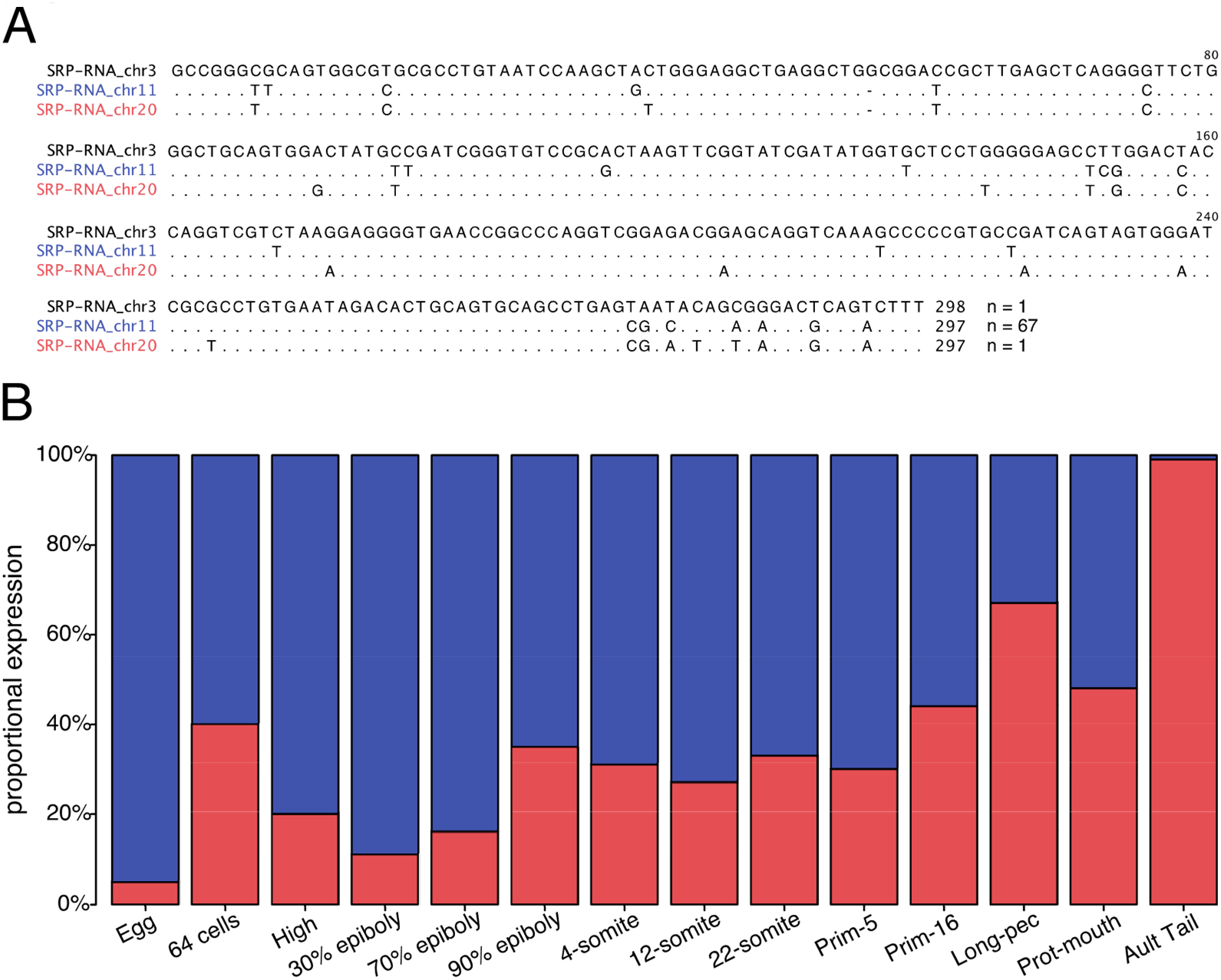
Maternal-type and somatic-type SRP-RNA variants. A: Sequence alignment of the zebrafish SRP?RNA gene variants. Identical nucleotides or gaps are indicated as dots and dashes, respectively. B: Bar plot showing the maternal-type (blue) and somatic-type (red) SRP-RNA proportional expression during zebrafish developmental stages (Supplemental Table ST3).

### Zebrafish RNA P maternal expression

To determine possible differential gene expression, we mapped the RNAseq reads from zebrafish eggs, a developmental embryonic series, and mature samples to the three rpph1 variant sequences (cf. M&M (Locati et al. 2017b), and Supplemental Table ST2).

The annotated rpph1 gene is the most prominently expressed variant in virtually all samples, in contrast to the similar copy, rpph1-sim, which is hardly ever expressed (Supplemental Table ST2). The dissimilar copy, rpph1-dis, is mainly expressed during oogenesis and embryogenesis, which is in line with it being the maternal-specific RNase P RNA gene. In contrast to most other somatic-specific non-coding RNAs, which are hardly present in egg, rpph1 transcripts are also quite prominently present in egg. Yet, it is important to emphasize that the analysis shows relative expression and the absolute expression of RNase P RNA is very low in egg and embryonic samples, as compared to low-proliferative tissues like brain and liver (Supplemental Table ST2). Additionally, an inspection of the mapped reads (data not shown) revealed other nucleotide differences as compared to the rpph1 reference sequences, suggesting the possible presence of one or more variants.

### Conclusion

There are three rpph1 variants in close proximity located on chromosome 2; the original rphh1 is expressed in all samples tested, one (similar) variant is hardly present, and one (dissimilar) variant is present virtually just in egg and during embryogenesis, which makes this the maternal-specific variant. However, all rrph1 expression levels in egg and embryonic samples are rather low compared to that of the ubiquitous rphh1 variant in somatic tissues. Given the numerous sequence differences between the two rpph1 variants, it is likely that most, if not all, protein binding sites are affected, which may have a profound effect on the RNase P RNP as a whole. For example, a study involving mutations in a RNase P protein in *drosophila* showed that these alterations induced complete sterility by triggering several DNA damage checkpoints in the oogenesis (Molla-Herman et al. 2015). It is impossible to assert which task(s) the maternal-type rpph1 has in the oogenesis and embryogenesis, other than it is related to a part of the maternal-specific translation system.

## SRP-RNA

### Background

The signal recognition particle (SRP) is a ribonucleoprotein responsible for the translocation of the mRNA-ribosome translation complex to the endoplasmic reticulum (ER). The SRP recognizes the ER signals sequence at the N-terminus of an integral membrane protein or secretory protein that is being synthesized. Direct binding of the SRP to the signal sequence peptide and indirect binding to ribosome stalls protein synthesis, after which the whole complex is guided to the ER. Binding to the SRP receptor in the ER membrane releases the SRP from the complex and translation resumes by synthesizing the protein through the ER membrane via a protein translocator (Walter and Blobel 1980; Lütcke 1995; Keenan et al. 2001; Akopian et al. 2013; Alberts et al. 2015).

The SRP consists of several proteins and a small non-coding RNA; SRP-RNA (also known as 7SL RNA), which is produced by RNA polymerase III (Walter and Blobel 1982; Leung and Brown 2010). This SRP-RNA forms the backbone of the SRP, is about 300 nucleotides long, and has a highly conserved secondary structure with a small (Alu) domain and a large (S) domain (Walter and Blobel 1983). The Alu domain is responsible for translation retardation and the S domain recognizes the signal sequence and also binds to the ER SRP receptor (Lütcke 1995; Wild et al. 2019).

### Zebrafish SRP-RNA genomic organization

In the zebrafish genome, SRP-RNA genes can be found on three chromosomes; 3, 11, and 20 with a standard BLAST analysis (Madden 2002). On chromosome 11, 67 complete copies are organized mainly in three gene clusters with a mutual homology of 91% and up. Both chromosome 3 and chromosome 20 contain one copy of the SRP-RNA gene with 93% and 94% homology to the SRP-RNA consensus sequence of the genes on chromosome 11 (Figure 3A, Supplemental Table ST3). The position of the gene copy on chromosome 20 is conspicuous, as is it approximately 10 kb downstream of the maternally expressed tRNA-LeuCAA cluster. The copy on chromosome 3 is located just 448 bp from the start of the *ER membrane protein complex subunit 10* gene.

### Zebrafish SRP-RNA maternal expression

The homology between the different SRP-RNA gene copies is relatively high, but there are sufficient sequence differences between the three chromosomal versions to identify them (Figure 4A). Mapping the RNAseq reads from egg, the embryonic development series, and adult samples to the three SRP-RNA variant sequences (cf. M&M (Locati et al. 2017b)) showed a clear differential expression (Figure 4B, Supplemental Table ST3). Transcripts from the SRP-RNA gene clusters on chromosome 11 are predominantly present in egg plus early embryogenesis and are during embryogenesis replaced by those from the single SRP-RNA gene on chromosome 3, which then becomes the almost exclusive variant expressed in adult tissues. The single copy on chromosome 20 is hardly expressed in any of the samples. The resulting expression patterns are highly similar to maternal-type rRNA (similar to 67 copies on chromosome 11,), somatic-type rRNA (similar to single copy on chromosome 3), and unexpressed-type rRNA (similar to single copy on chromosome 20) (Locati et al. 2017a, 2017b).

### Conclusion

The SRP-RNA seems to be a part of the maternal-specific translation system as there is clear differential expression between the gene copies. Intriguingly, the SRP-RNA genomic organization seems to follow the typical trend that maternal variants are present in large numbers and in clusters, whereas the somatic variants are often solitary gene copies.

The sequence differences between the maternal chromosome 11 SRP-RNA copies and the somatic (chromosome 3) variant (~7%) are both in the Alu domain, as well as in the S domain. As several SRP proteins bind directly to the SRP-RNA, the sequence differences might affect binding of them, so the maternal-type variant SRP might end up being significantly different from the somatic-type variant.

Altogether, zebrafish has a maternal-specific SRP to guide the mRNA-ribosome complex to the ER. As to why also this part of the translation system is present in a maternal-type variant remains elusive for now, although it seems possible that is a, yet unidentified element of the ribosome filter hypothesis (Mauro VP, 2002, 2007, and 2016), which seems related at least to the 45S rRNA maternal- and somatic types (Locati et al. 2017b). In any case, as virtually no research is done into the zebrafish SRP-RNA, the discovered dual system provided an excellent opportunity to study this.

## Conclusion

In this manuscript, we presented the preliminary results of our initial gene-expression analyses of the non-coding RNA types involved in the zebrafish translation system, other than we previously reported on (Locati et al. 2017a, 2017b, 2018; Pagano et al. 2019b, 2019a). Alike rRNA (Locati et al. 2017a, 2017b), snoRNA (Pagano et al. 2019b), and snRNA (Pagano et al. 2019a), the investigated tRNA, RNase-P and SRP-RNA all seem to have a maternal-specific variant. Together, a complete maternal-specific translation system emerges, which is dedicated to oogenesis and (early) embryogenesis in zebrafish. Especially the existence of a maternal-specific SRP-RNA points to a complete system, as it targets the initial part of the produced protein. Besides the maternal-specific expression profiles of the RNA-processing snoRNA (Pagano et al. 2019b) and RNase P (both of which are indirect non-coding RNA actors in the translation system cf. Figure 1), all translation-involved non-coding RNA types show rather absolute expression in egg, virtual absent expression in adult tissue, and decreasing expression during embryogenesis.

Putting the findings of the here presented studies in context with our previous finding of non-coding RNAs that are involved in the cellular translation system, leads to a number of intriguing reflections.

### Maternal-type proteins

All translation-involved non-coding RNAs function as ribonucleoprotein particles (RNPs), but often depend on RNA-RNA interactions. The nucleotide differences between maternal-type and somatic-type RNA variants are mostly located at the protein binding sites of the implicated non-coding RNAs. This raises the question whether the translation RNP-associated proteins, for instance between the maternal-type and somatic-type ribozymes, are also dissimilar. A quick search did not show maternal-specific expression of the protein-coding genes of these RNPs, however, as these genes contain introns, possible different proteins could be orchestrated via differential splicing.

### An “unexpressed-type” translation system

One of the first steps in the performed analyses of each non-coding RNA type was to characterize their complete genomic presence. Once we identified all the complete gene copies of a RNA type, their expression patterns in zebrafish development were determined. Besides the expressed maternal-type and somatic type RNAs, we discovered several apparently functional genes that were not, or extremely low expressed. For this reason, we labeled these gene copies as “unexpressed”. Unexpressed-type gene copies have been identified for: rRNAs, snoRNAs, snRNAs, RNase and SRP-RNA (for tRNAs we were unable to perform a complete genomic characterization). Because we assume that these variants have to be expressed somewhere, several research attempts were made to find transcripts of these unexpressed-type non-coding RNAs. As we speculated that this might actually be a paternal-type variant, we analyzed zebrafish testis samples, as well as various other tissues (unpublished results). The unexpressed-type variant turned out to be not expressed in testis and the results of the other tissues were inconclusive given the overwhelming presence of maternal- and somatic-type transcripts in the RNAseq data. It may be that this particular variant is only expressed in certain low-frequency cell types, for instance stem cells, or alternatively it is only expressed in response to stimuli such as stress, hormones, or temperature. In any case, determining where this unexpressed translation system variant is expressed will be extremely informative for unraveling the relevance of the involved (sequence) differences between all three translation systems.

### Genomic organization

One of the most obvious distinctions between the maternal-type and somatic-type RNA variants is their particular genomic organization. There are almost always many maternal-specific gene copies, whereas of the somatic-type gene copies there are often just a few. The maternal loci have a hallmark organization of repeated and clustered genes with a preference for chromosome 4 (Table 1). Notably, this chromosome is identified as a gender chromosome in zebrafish (Anderson et al. 2012; Nagabhushana and Mishra 2016). The somatic-type gene copies are virtually always solitarily organized. For 5S rRNA we could show a connection between a specific retrotransposon family and the maternal-type tandem arrays (Locati et al. 2017a). Given the absence of complete retrotransposons in the somatic-type 5S rRNA locus, we suggested an involvement of this specific retrotransposon family in achieving and maintaining the differential copy number of the two types of 5S rDNA loci. Although we did not fully analyze this phenomenon in the other translation-involved non-coding RNA loci, it was clear that several of them showed similar retrotransposon hallmarks such as; partial gene sequences and genomic spaces in otherwise tightly-packed tandem arrays (unpublished results). It appears as if there is a specific retrotransposon family involved in almost each of the mentioned non-codon RNAs. This could hint to a universal mechanism to maintain a proper number of functional gene copies (Kojima 2003; Dupuis-Sandoval et al. 2015). In any case, identifying the exact retrotransposons associated with the relevant non-coding RNA loci might elucidate the genome organization of these RNAs, as well as their possible non-canonical functions (Buzdin et al. 2007).

### Proliferation-specific translation system

One of the nagging issues for all of the non-coding RNA research is the lingering question as to why there is an apparent need in zebrafish for a dual (or even triple) different translation system. In an early stage of our research, we named the newly-discovered alternative system “maternal-type” due to its transcription origin in oocytes. As a logic consequence, the other translation system became “somatic-type”. However, this decision came back to haunt us, when we much later discovered that testis tissue, presumably (immature) sperm cells, contain a considerable amount of maternal-type translation-involved non-coding RNA. Another option could be that our maternal-type translation system actually is related to stem cells in general, not just the totipotent fertilized egg cell. Recently there have been several reports on alternative snRNAs specifically expressed in stem cells (Vazquez-Arango et al. 2016; Kim et al. 2017; Schmidt et al. 2018). In line with these observations is the report on the existence of two distinct human translation programs that operate during either cell proliferation or differentiation (Gingold et al. 2014). This aligns with a large body of research that implicates (variant) nucleolus components in cancer cells (Derenzini et al. 1998; Lo et al. 2006; Jorjani et al. 2016). Given that the first phase in zebrafish embryogenesis of about eight hours is primarily dedicated to proliferation, after which is switch is made to mainly differentiation, it could well be that our maternal-type translation system is in reality a proliferation-specific translation system.

## SUPPLEMENTAL TABELS

ST1.xlsx: tRNA

ST2.xlsx: RNase

ST3.xlsx: SRP

## Supporting information

Supplemental Tabel ST1

Supplemental Tabel ST2

Supplemental Tabel ST3

